# Identification of human vaccinees that possess antibodies targeting the egg-adapted hemagglutinin receptor binding site of an H1N1 influenza vaccine strain

**DOI:** 10.1101/249615

**Authors:** Tyler A. Garretson, Joshua G. Petrie, Emily T. Martin, Arnold S. Monto, Scott E. Hensley

## Abstract

Human influenza viruses passaged in eggs often acquire mutations in the hemagglutinin (HA) receptor binding site (RBS). To determine if egg-adapted H1N1 vaccines commonly elicit antibodies targeting the egg-adapted RBS of HA, we completed hemagglutinin-inhibition assays with A/California/7/2009 HA and egg-adapted A/California/7/2009-X-179A HA using sera collected from 159 humans vaccinated with seasonal influenza vaccines during the 2015-16 season. We found that ~5% of participants had ≥4-fold higher antibody titers to the egg-adapted viral strain compared to wild type viral strain. We used reverse-genetics to demonstrate that a single egg-adapted HA RBS mutation (Q226R) was responsible for this phenotype.

## Introduction

Influenza viruses attach to cells through specific interactions between the viral hemagglutinin (HA) protein and terminally-linked sialic acids on target cells. Human influenza viruses possess a conserved HA receptor binding site (RBS) that interacts efficiently with α-2,6-linked sialic acids, whereas the HA RBS of avian influenza strains primarily interacts with α-2,3-linked sialic acids [1]. Most human influenza vaccine antigens are prepared from viruses grown in fertilized chicken eggs. Human influenza viruses grown in eggs often acquire mutations in or near the HA RBS that increase binding to α-2,3-linked sialic acids, and some of these mutations lead to large antigenic changes [2]. For example, the 2016-17 H3N2 egg-grown vaccine was antigenically mismatched compared to circulating H3N2 strains due to a T160K HA mutation that arose during egg passage [3]. In this case, the egg-adapted T160K HA mutation was located in a classic antigenic site adjacent to the RBS [3].

Recent studies have identified antibodies with long CDR3 domains that essentially act like sialic acid mimics that make physical contact with HA RBS residues [4, 5]. There is considerable interest in developing vaccines that elicit these types of antibodies since they are able to neutralize a wide range of different influenza virus strains. It is unclear if vaccine strains with egg-adapted RBSs are able to elicit these broadly reactive antibodies, given that most egg-grown vaccine strains possess RBS mutations that facilitate growth in eggs. In a landmark study, Raymond and colleagues isolated monoclonal antibodies from vaccinated humans that bind to the egg-adapted RBS of H1N1 but not to circulating H1N1 viral strains [6]. These antibodies bind to egg-grown H1N1 viral strains that utilize α-2,3-linked sialic acids but not to viral strains that actually circulate in humans that utilize α-2,6-linked sialic acids [6]. It is unknown if these types of antibodies are commonly elicited by egg-adapted H1N1 vaccine strains. To address this, we completed hemagglutination-inhibition (HAI) assays with ‘wild-type’ H1N1 HA (A/California/7/2009) and ‘egg-adapted’ H1N1 HA (A/California/7/2009-X-179A) using sera collected from 159 individuals pre- and post-vaccination with the egg-adapted 2015-2016 seasonal influenza vaccine.

## Methods

Prior to the 2015-2016 influenza season, individuals were enrolled in the University of Michigan Household Influenza Vaccine Effectiveness (HIVE) study, as previously described [7, 8]. Households were defined as having ≥4 members, of which ≥2 were children under age 18. Over the course of the 2015-16 influenza season, nasal and throat swab samples were collected from participants that displayed symptoms of acute respiratory illnesses and these samples were tested for influenza virus by real-time reverse-transcription polymerase chain reaction (RT-PCR). For this study, we analyzed sera from participants that received a seasonal influenza vaccine (97% received Sanofi Pasteur vaccine, 1% received GSK vaccine, 2% vaccine type unknown). Serum samples were collected from participants ages ≥13 years at the time of enrollment and also ≥14 days following vaccination. Informed consent was obtained and the study was approved by the University of Michigan Medical School Institutional Review Board.

HAI assays were performed using de-identified sera collected pre- and post-vaccination from 159 individuals with the approval of the University of Pennsylvania Institutional Review Board. Sera were pre-treated with receptor-destroying enzyme for 2 hours at 37°C and inactivated for 30 minutes at 55°C. Sera were also absorbed with 10% red blood cell solution for 1 hour at 4°C prior to completing HAI assay. We used influenza virus-like particles (VLPs) for the HAI assays in this study as previously described [9], since it is difficult to grow human H1N1 viruses without adaptive mutations. We used VLPs that expressed the ‘wild-type’ A/California/7/2009 H1N1 HA or the egg-adapted A/California/7/2009-X-179A H1N1 HA. The VLPs for these experiments possessed a neuraminidase (A/duck/Alberta/300/77) that most humans have not been exposed to previously. We constructed our VLPs in this manner to avoid potential complications from neuraminidase-reactive antibodies. We also included 20nM of oseltamavir in our HAI assays to prevent neuraminidase binding of red blood cells [10]. The A/California/7/2009 wild-type HA and the A/California/7/2009-X-179A HA in our VLP constructs differ at 2 residues; the X-179A strain possesses a glutamine to arginine mutation at position 226 (Q226R) and from lysine to threonine at position 212 (K212T). For some experiments we completed additional HAI assays using VLPs that expressed A/California/7/2009 HAs that were engineered to possess only the Q226R mutation that is in the RBS.

## Results

We completed two independent HAI assays using sera from 159 individuals collected prior to and ≥14 days after vaccination. Vaccinated subjects ranged in age from 13 to 76 years old: 22 (14%) were ≤18 years old and 22 (14%) were ≥50 years old. We completed HAI assays with VLPs that possessed the A/California/7/2009 H1N1 wild-type HA or the A/California/7/2009-X-179A HA. The wild-type HA differs from the egg-adapted X-179A HA at residues 212 (K212T) and 226 (Q226R) (Figure 1). Residue 212 is somewhat buried within the trimer interface, whereas residue 226 is located directly in the RBS (Figure 1) and is known to impact receptor binding [11]. HAI titers from two independent experiments are shown for all participants in Supplemental Table 1.

**Figure 1.**
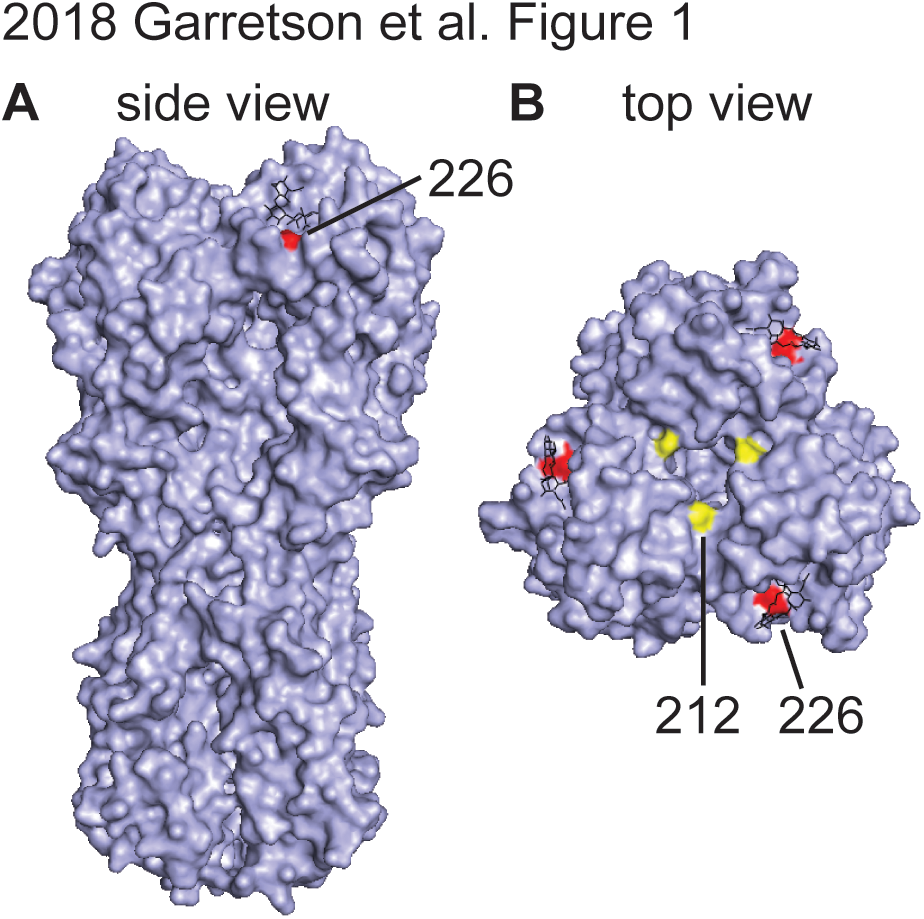
Differences between WT and X-179A HAs. A side view (A) and top view (B) of the H1 trimer are shown (PDB ID code 3UBN). Residue 212 is shown in yellow and residue 226 is shown in red. The glycan receptor is shown in black.

We identified 8 participants (~5% of participants that we tested) that had antibody titers that were ≥4-fold higher to the egg-adapted X-179A HA compared to the wild-type HA following vaccination in 2 independent HAI experiments (Supplemental Table 1). Some of these individuals possessed antibodies that reacted to the X-179A HA better than the WT HA prior to vaccination, which is likely the case because the X-179A vaccine strain has been utilized in the human population since 2009. However, some individuals clearly seroconverted against the X-179A HA but not the WT HA following vaccination in this study. For example, participant #81’s HAI titer against X-179A HA rose from a value of 15 pre-vaccination to a value of 100 post-vaccination but HAI titers in this individual against WT HA were nearly the same pre- and post-vaccination (pre-vaccination titer of 10 and post-vaccination titer of 25) (Supplemental Table 1; average HAI titer from 2 experiments described in text). Similarly, participant #147 was HAI negative against both HAs prior to vaccination and had a much higher HAI titer against X-179A HA (HAI titer of 70) compared to WT HA (HAI titer of 10) following vaccination (Supplemental Table 1; average HAI titer from 2 experiments described in text). These data indicate that the X-179A egg-adapted vaccine strain elicits and/or reinforces antibody responses that recognize the X-179A egg-adapted H1N1 HA more efficiently than the WT H1N1 HA in some individuals.

To confirm that the difference in WT HA versus X-179A HA reactivity was due to RBS differences, we completed additional HAI experiments using A/California/7/2009 HAs that were engineered to possess only the Q226R RBS HA mutation. HAI titers were lower using the WT HA compared to the HA with the Q226R mutation (Figure 2), indicating that the 8 individuals that we identified in our study possessed antibodies that targeted the egg-adapted RBS of the X-179A H1N1 vaccine strain.

**Figure 2.**
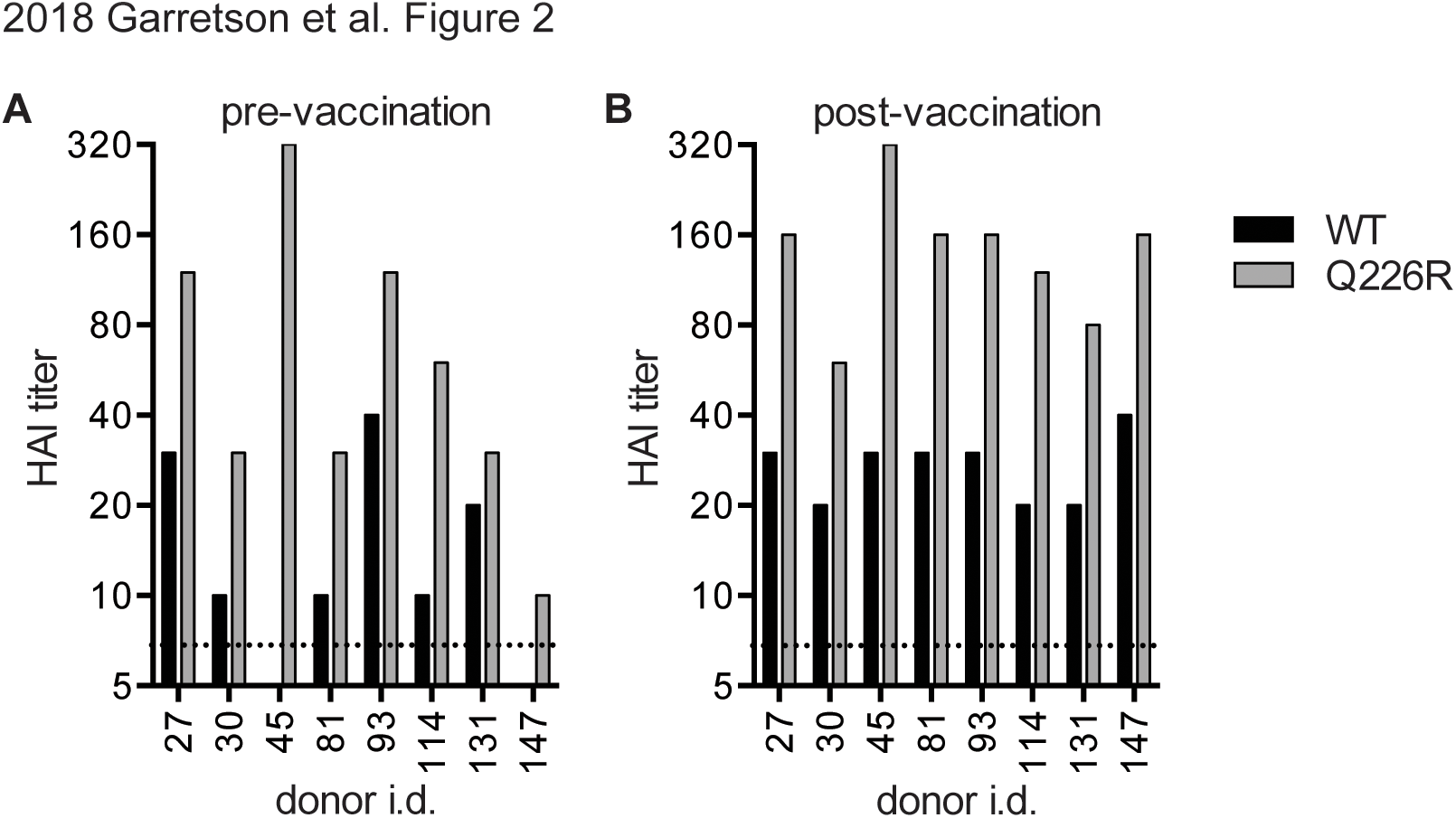
Identification of individuals that possess antibodies that target the RBS of X-179A following vaccination. HAI assays using VLPs with WT HA and HA engineered to possess the Q226R mutation were completed using sera from participants that had ≥4-fold higher HAI titers to the egg-adapted X-179A HA compared to the wild-type HA as determined in Supplemental Table 1. Sera collected prior to (A) and after (B) vaccination were tested. Data are representative of 2 independent experiments.

Seven out of the 159 vaccinated individuals in this study had PCR-confirmed H1N1 infection during the course of the 2015-16 season. All 7 of these individuals possessed low antibody titers against the WT H1N1 strain after vaccination (Supplemental Table 1). Importantly, one of the H1N1-infected participants (#131) had a higher HAI titer to the X-179A HA compared to the WT HA following vaccination, albeit titers to both HAs were low in this individual. From these studies, we conclude that some individuals vaccinated with the egg-adapted X-179A vaccine strain produce antibodies that recognize egg-adapted epitopes in the HA RBS and we speculate that this might contribute to reduced vaccine effectiveness.

## Discussion

Human influenza viruses typically acquire mutations in and around the HA RBS that enhance virus binding to α-2,3-linked sialic acids when grown in fertilized chicken eggs. Essentially all human influenza vaccines propagated in eggs possess egg-adapted HA mutations. Sometimes these mutations are located in classical antibody binding sites that lead to substantial vaccine mismatches, as was the case during the 2016-2017 H3N2-dominated influenza season [3]. More commonly, egg-grown vaccine strains possess adaptive mutations ‘buried’ in the RBS, and it has been historically thought that these mutations are not as antigenically important. However, it is becoming clear that some human antibodies with long CDR3 domains physically make contact with residues deep within the RBS [4, 5], and a recent study isolated monoclonal antibodies from a vaccinated human that could bind to HAs with an egg-adapted RBS but not to non-egg adapted HAs that were present in viruses that circulated in humans [6]. Here, we set out to determine how common these types of antibodies are in vaccinated humans and if they are associated with vaccine failure.

We found that ~5% of individuals vaccinated with the 2015-2016 seasonal influenza vaccine possessed antibodies that recognized the X-179A H1N1 HA ≥4-fold more efficiently compared to the WT H1N1 HA. Given that >140 million people receive a seasonal influenza vaccine in the U.S. each year [12], these data indicate that a large number of individuals (>7 million!) possess antibodies that preferentially recognize the egg-adapted HA RBS of H1N1 rather than the HA of H1N1 viruses that circulate in humans. For our study, we focused on individuals that had ≥4-fold differences in HAI titers using the different HAs, and this conservative fold difference cutoff likely underestimates the number of individuals that possess antibodies that bind to the egg-adapted H1N1 strain but not the circulating H1N1 strain. It is notable that participants in our study come from a population with historically high annual vaccine uptake rates, and therefore they are likely to have had multiple previous influenza vaccines, as well as a low burden of chronic diseases that may impair antibody responses [13]. Consistent with this, several participants in our study possessed antibodies that recognized the X-179A HA more effectively than the WT HA prior to vaccination in 2015-2016. Since the egg-adapted H1N1 component of seasonal influenza vaccines remained unchanged between 2010-2016, it is possible that these participants developed this phenotype during an earlier vaccination and that these antibody responses were continually boosted by subsequent immunizations.

While most studies of egg-adaptation have focused on H3N2 viruses, our study clearly demonstrates that egg-adaptations in the HA RBS of H1N1 viruses can also be problematic. Antibodies targeting the RBS of HA have the potential to be broadly neutralizing if they are not restricted to binding only egg-adapted HAs [14, 15]. Our current reliance on eggs for the production of most seasonal influenza vaccines disfavors the generation of these types of broadly reactive antibodies. It is important to continue to develop new systems to prepare influenza vaccine antigens that are not dependent on viral growth in eggs, such as baculovirus and mammalian cell-based systems. Future studies should address if antigens produced in these alternative systems are more effective at eliciting antibodies that target the RBS of HAs that are actually present in human influenza virus strains.

